# Encoding time in neural dynamic regimes with distinct computational tradeoffs

**DOI:** 10.1101/2021.07.15.452447

**Authors:** Shanglin Zhou, Sotiris C. Masmanidis, Dean V. Buonomano

## Abstract

Converging evidence suggests the brain encodes time in time-varying patterns of neural activity, including neural sequences, ramping activity, and complex dynamics. Temporal tasks that require producing the same time-dependent output patterns may have distinct computational requirements in regard to the need to exhibit temporal scaling or generalize to novel contexts. It is not known how neural circuits can both encode time and satisfy distinct computational and generalization requirements, it is also not known whether similar patterns of neural activity at the population level can emerge from distinctly different network configurations. To begin to answer these questions, we trained RNNs on two timing tasks based on behavioral studies. The tasks had different input structures but required producing identically timed output patterns. Using a novel framework we quantified whether RNNs encoded two intervals using either of three different timing strategies: scaling, absolute, or stimulus-specific dynamics. We found that similar neural dynamics for single intervals were associated with fundamentally different encoding strategies and network configurations. Critically, some regimes were better suited for generalization, categorical timing, or robustness to noise. Further analysis revealed different connection patterns underlying the different encoding strategies. Our results predict that apparently similar neural dynamic regimes at the population level can be produced through fundamentally different mechanisms—e.g., in regard to network connectivity and the role of excitatory and inhibitory neurons. We also predict that the task structure used in different experimental studies accounts for some of the experimentally observed variability in how networks encode time.

**Author summary:** The ability to tell time and anticipate when external events will occur are among the most fundamental computations the brain performs. Converging evidence suggests the brain encodes time through changing patterns of neural activity. Different temporal tasks, however, have distinct computational requirements, such as the need to flexibly scale temporal patterns or generalize to novel inputs. To understand how networks can encode time and satisfy different computational requirements we trained recurrent neural networks (RNNs) on two timing tasks that have previously been used in behavioral studies. Both tasks required producing identically timed output patterns. Using a novel framework to quantify how networks encode different intervals, we found that similar patterns of neural activity—neural sequences—were associated with fundamentally different underlying mechanisms, including the connectivity patterns of the RNNs. Critically, depending on the task the RNNs were trained on, they were better suited for generalization, categorical timing, or robustness to noise. Our results predict that similar patterns of neural activity can be produced by distinct RNN configurations, which in turn have fundamentally different computational tradeoffs. Our results also predict that differences in task structure account for some of the experimentally observed variability in how networks encode time.

## Introduction

The ability to predict when external events will occur, and to detect temporal regularities in the environment, are among the most fundamental computations the brain performs [1–5]. Thus, the brain must have a rich repertoire of mechanisms to tell time and perform temporal computations. Indeed, converging experimental and computational evidence indicates that a wide range of different brain areas encode time through dynamically changing patterns of neural activity [1, 6–10]. These patterns can take the form of monotonic ramping of the firing rates of neurons, or so-called population clocks that can take the form of neural sequences or complex patterns of neural activity [1, 11].

Experimental and computational analyses of the different neural encoding schemes for the representation of time have focused primarily on the discrimination and production of isolated intervals or durations. However, the computational requirements for processing temporal information go far beyond merely using a timer to discriminate or produce a single duration or interval. Some forms of temporal processing require the ability to smoothly scale a time-varying motor pattern. For example, the ability to play a song on the piano at different tempos, or catch a ball thrown at different speeds, requires that the underlying patterns of neural activity unfold at different speeds [12–15]. Indeed, some tasks in animal studies explicitly require animals to exhibit temporal scaling: depending on context cues or training blocks animals must temporally scale their motor response [14, 16–18]. In contrast, other timing tasks are categorical in nature, for example in the language domain phrasal boundaries are based in part on a categorical boundary of the pause between phonemes—e.g., *great eyes* x *gray ties* [19, 20], similarly, in the motor domain, the distinction between a double-click and two single clicks of a computer mouse is categorical. Furthermore, in both the human and animal literature standard temporal bisection tasks require subjects to make a two-alternative forced-choice categorical judgment regarding whether a stimulus was short or long [21, 22].

We propose that depending on task structure and requirements there are computational tradeoffs between different encoding schemes and patterns of neural activity the brain may use to encode time. Consider a task in which an animal has to produce two intervals—e.g., in response to two different sensory cues. Generally speaking, three encoding schemes could allow the same network to produce these two different intervals: absolute timing, temporal scaling, and stimulus-specific timing. Under *absolute* timing the neurons would respond at the same moments in time during both the production of short and long intervals but additional neurons would be active during the long interval; in a *temporal scaling* scheme neurons encode the same relative time during both short and long intervals; and in a stimulus-specific code, there would be unrelated patterns for each interval (e.g., entirely different neural sequences for the short and long interval). These different schemes possess specific computational tradeoffs regarding their suitability for temporal scaling versus categorical timing.

To date, a large diversity of neural signatures for time—including scaling, absolute timing, and stimulus-specific timing—have been observed during tasks that require animals to discriminate or produce multiple intervals [14, 16–18, 23–30]. Here we propose that some of this diversity is driven by task structure, and examine whether task structure influences the way recurrent neural networks may encode time. To address this hypothesis we trained RNNs on two tasks with identical output motor requirements and characterized how the networks encode time and generalize to novel stimuli. Our results establish that subtle differences in task structure lead to neural dynamic regimes that are better suited for temporal scaling or categorical timing.

## Results

To begin to understand how task structure might shape how time is encoded in neural networks, we trained recurrent neural network models (RNNs) on one of two tasks inspired by previous experimental studies[14, 23]. The RNNs were based on firing rate units with distinct population of excitatory (80%) and inhibitory (20%) units. We will refer to the tasks as the 2-Context (Fig 1**A**) and 2-Stimulus (Fig 1**B**) tasks—critically, the timed motor outputs were identical in both tasks, requiring the production of either a short or long response. In the 2-Context task, the *go* cue (500 ms) indicated the onset of the trial (t=0), and the analog level of a continuous context input signaled whether a trial is short or long[e.g., 14]. In the 2-Stimulus task, the short and long interval trials were cued by two distinct transient inputs [23]. In both cases, the short and long intervals consisted of a ramp-up of the output unit starting at the interval midpoint—a function that approximates the behavioral response rate of animals trained to correctly time their movements [23].

**Fig. 1.**
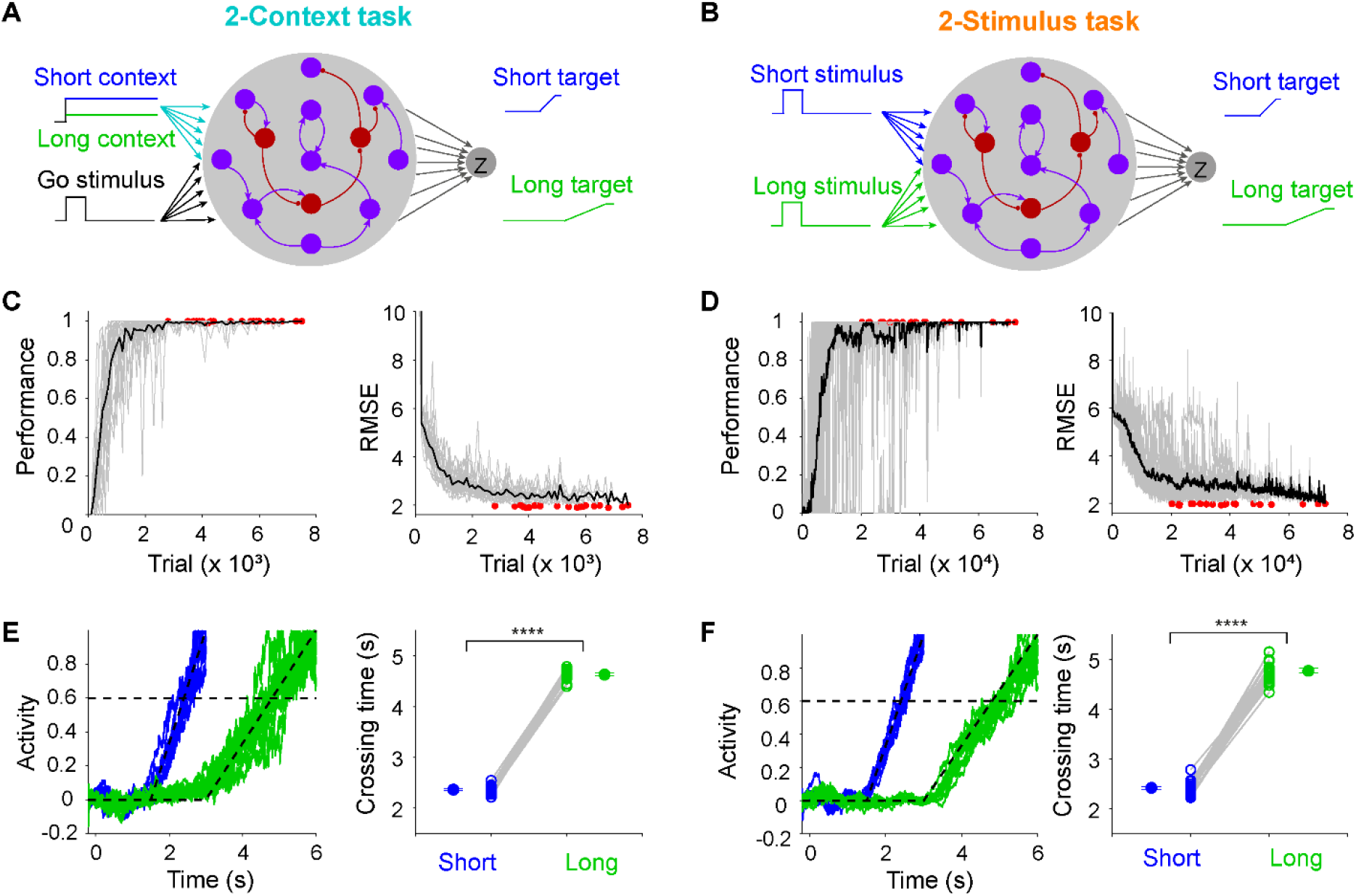
RNNs were trained on one of two timing tasks, both of which required producing the same timed output patterns. (**A**) Schematic of the 2-Context task. Each RNN was composed of 200 units—80% excitatory units (purple) and 20% inhibitory units (dark red)—and received a *go* and a *context* input. The context level signals the interval length to be produced: high = long (6 s, blue), low = short (3 s, green). (**B**) Schematic of the 2-Stimulus task. The same RNN was used in both tasks, except that the short- and long-interval was cued by two different inputs that were transiently activated. (**C**) Learning curve for the performance of 20 RNNs trained on the 2-Context task. Percentage of trials in which the timing of the output unit met criteria (left) and the RMSE between the output and target (right). Gray traces represent results of each RNN, red dots denote the end of training for a given RNN, and the black trace represents the mean performance. (**D**) Same as in (**C**) but for the 2-Stimulus task. (**E**) Output traces across ten short (blue) and long (green) trials from an RNN trained on 2-Context task (left). Mean crossing times for long interval is significantly higher than that for short interval (right, *n* = 20 simulations, paired t test, *t_19_* = 77.70, *P* < 0.0001). Dashed lines denote the targets and threshold. (**F**) Same as (**E**) but for 2-Stimulus task (*n* = 20 simulations, paired t test, *t_19_* = 45.79, *P* < 0.0001).

Performance was quantified by the ratio of correctly timed trials (see Methods) and RMSE. RNNs trained on both tasks learned to produce the same appropriately timed motor output (Fig 1**C-F**). RNNs trained on the 2-Context task, however, required fewer training trials to reach the same performance level (n = 20 simulations, two-sample two-sided t-test, t_38_ = 9.75, P < 0.0001).

### Generalization to novel intervals

Having shown that RNNs can produce the same temporal output patterns when trained on two similar tasks, we next asked a key question: are there significant computational tradeoffs between how both RNNs trained on the different tasks perform in response to novel conditions? To answer this question we examined generalization to untrained input conditions. To test the generalization in the 2-Context task we varied the amplitude of the context cue between the range of the trained values (0.75=short; 0.25=long). Interestingly the network exhibited fairly smooth generalization—i.e., in response to intermediate context levels it produced intermediate motor intervals (Fig 2**A**)—a finding consistent with previous computational studies [12, 14]. To test generalization in the 2-Stimulus task we mixed the ratio of activation of the two stimulus cues—during training [1, 0] corresponded to short and [0, 1] to long, during testing an intermediary 50/50 mixed input corresponded to [0.5 0.5]. In contrast to the 2-Context task, the RNNs trained on the 2-Stimulus task did not generalize, but the RNNs did not exhibit catastrophic degradation or behave randomly. Rather, the RNNs expressed categorical timing: the output intervals clustered near the short or long intervals (Fig 2**B**), essentially exhibiting a winner-take-all behavior.

**Fig. 2.**
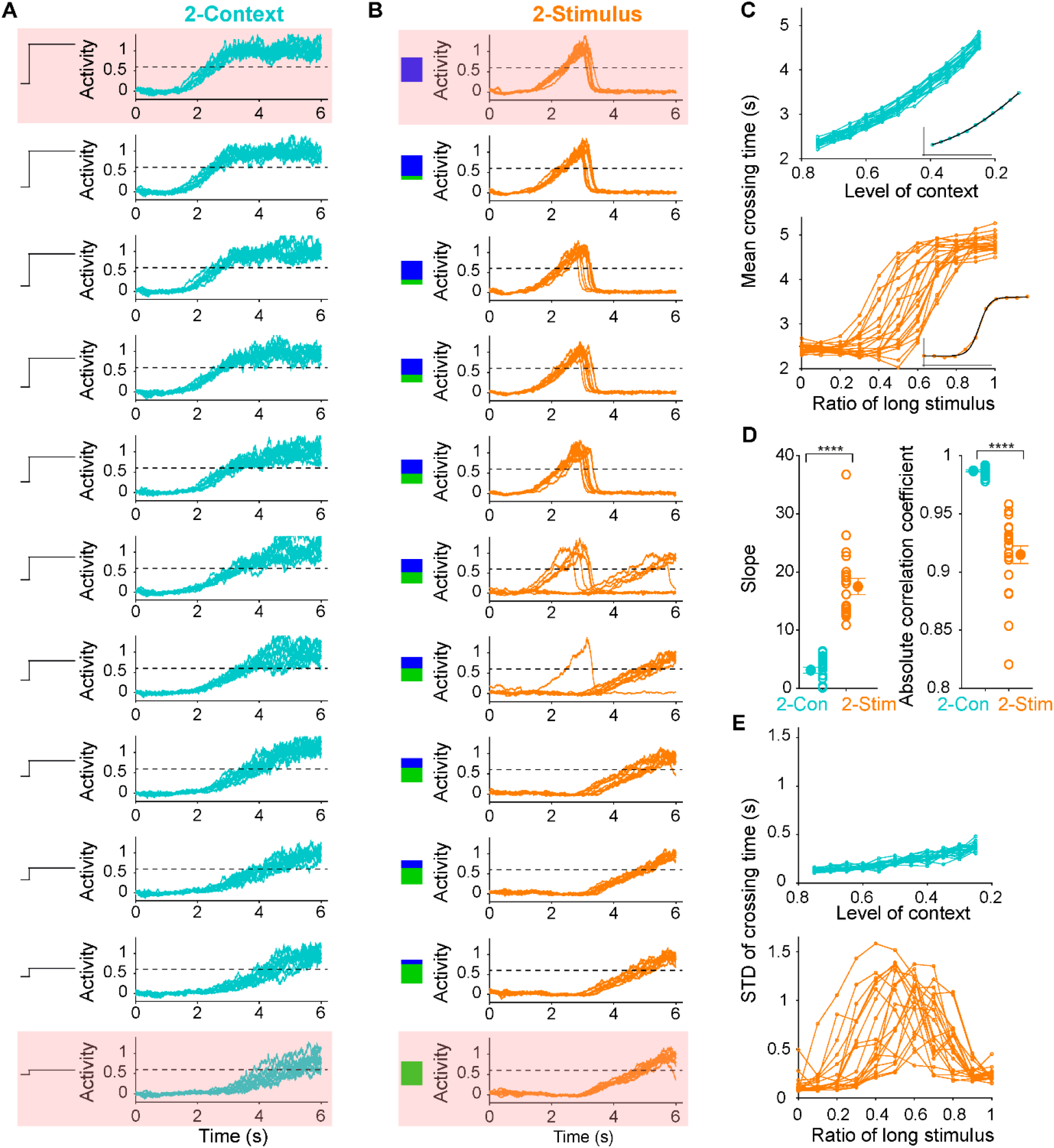
RNNs trained on the 2-Context task exhibited smooth generalization to novel intervals, while RNNs trained on the 2-Stimulus task exhibited categorical timing. (**A**) Output traces of an RNN trained on the 2-Context task across different context input levels. Dashed-black lines denote the output threshold used to quantify timing. Pink squares denote the trained conditions. (**B**) Similar to (**A**) but for the 2-Stimuls task. The blue and green squares represent the ratio of activation of the two input units. (**C**) Plots of the mean crossing time for each RNN across input conditions for the 2-Context (top) and 2-Stimulus (bottom) tasks. Insets, examples of the sigmoid-function fits for a single RNN (black). (**D**) Left, mean slope of the sigmoid fits for 2-Stimulus task is significantly higher than that for the 2-Context task (*n* = 20 simulations for each, two-sided t test, *t_38_* = 9.69, *P* < 0.0001). Right, correlation coefficient between mean crossing times and input conditions for 2-Context task is significantly higher than that for the 2-Stimulus task (*n* = 20 simulations for each, two-sided t test on Fisher-transformed values, *t_38_* = 17.39, *P* < 0.0001). (**E**) Standard deviations of the crossing times for each RNN in the 2-Context (top) and 2-Stimulus (bottom) tasks, as a function of input conditions.

To quantify these generalization patterns we measured the slope of a sigmoid fit between input levels and output intervals, as well as the correlation between them (Fig 2**C, D**, see Methods). The slope of the sigmoid was significantly less in the 2-Context fits—indicating a quasi-linear relationship between context input level and produced intervals. The sigmoid slope was significantly higher in the 2-Stimulus task, consistent with the prototypical sigmoidal signature of categorical discrimination (Fig 2**D**, left panel). Similarly, the Pearson correlation coefficients further supported the observation that the input-interval relationship was much more linear in the 2-Context task compared to the 2-Stimulus task (Fig 2**D**, right panel).

In addition to the above accuracy measures, we also quantified the precision of timing across the different generalization conditions, as the standard deviation of the crossing time of each trial (Fig 2**E**). The precision for the 2-Context task was high (low standard deviation) for all the stimulus conditions. In contrast, in the middle range for the 2-Stimulus task precision was very low. This was mainly due to categorical timing, i.e., in some stimulus conditions, the motor output would randomly be attracted towards the short or long interval. Taken together, RNNs trained for the 2-Context task were far superior at generalizing to novel intervals in terms of both timing accuracy and precision, however, the RNNs trained on the 2-Stimulus task were well suited for categorical timing.

### Potential dynamic regimes underlying the encoding of multiple intervals

Converging experimental and theoretical evidence indicates that a broad range of neural dynamic regimes encode time. But to date, these different regimes have not been contrasted in terms of their ability to encode multiple intervals and lead to generalization or categorical timing, or robustness to noise. Here we examine three broad potential strategies for the encoding of two intervals: scaling, absolute, and stimulus-specific codes. To illustrate these three strategies we consider how a network of neurons could encode both a short (3 s) and long (6 s) intervals (Fig 3)—note that while we use neural sequences to contrast the three encoding schemes, the same classification applies to other codes for time, including ramping activity. In a temporal scaling strategy (Fig 3**A**), the dynamics of each unit for the short interval is linearly scaled in time to produce the long interval (Fig 3**B**), which at the level of single units leads to two overlapping curves (Fig 3**C**). Similarly, when the neural trajectories of the entire population are projected into a low-dimensional space by principal component analysis the trajectories are also overlapping (Fig 3**D**). Under an absolute encoding strategy (Fig 3, middle panels) the temporal profile of each unit during the short interval does not change during the long interval. The long interval simply relies on recruiting additional neurons that have later temporal fields. Thus in PCA space, the curves for the short interval matched the first half of that for the long interval. In a stimulus-specific strategy (Fig 3, right panels), the temporal profile of each neuron is essentially uncorrelated during the short and long intervals. Thus in PCA space, the trajectories of the neural patterns of activity produced during the short and long intervals are distinct from one another.

**Fig. 3.**
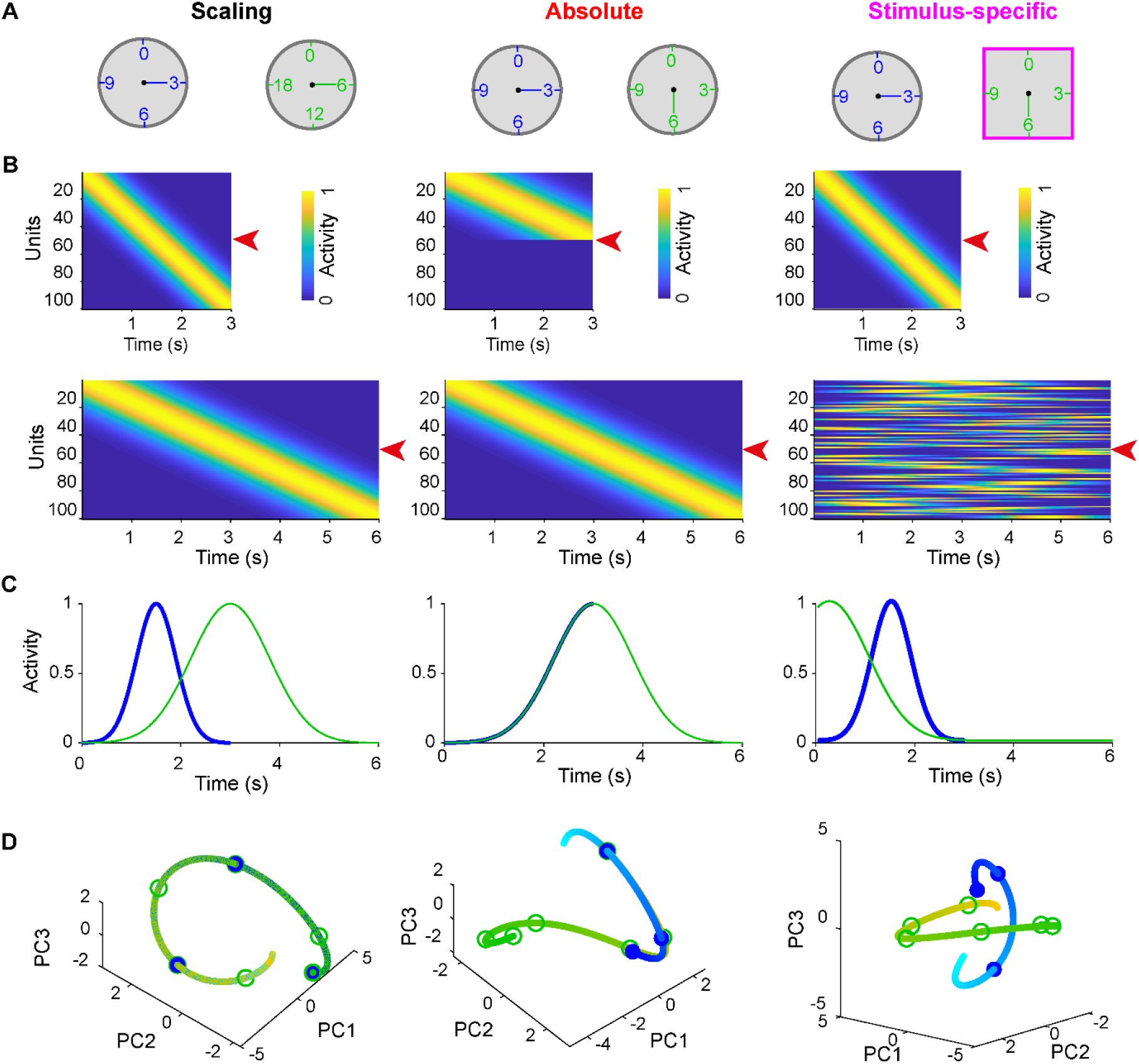
Three strategies for the encoding of two intervals by the same group of neurons. (**A**) Schematic of three potential strategies for timing two intervals: scaling, absolute, and stimulus-specific from left to right. (**B**) Prototypical dynamics for each of the encoding schems for a population of units during production of the short (top) and long (bottom) intervals. (**C**) Activity traces of the units denoted by the red arrows in (**B**) for short (blue) and long (green) interval. (**D**) Trajectories of three PCA components for short (cyan-blue) and long (yellow-green) interval for the corresponding population dynamics. The gradient colors (from the light to the dark) denote the flow of time. Circles denote the time points of the 1th, 2nd, 3rd, 4th, 5th, and 6th seconds.

Importantly, these encoding strategies are not necessarily mutually exclusive within a population of neurons. A network could use mixed encoding strategies in which different neurons are best described as scaling from one interval to another, while others encode absolute time. It is also possible that the dynamics of a given unit exhibit an absolute code early in a trial followed by scaling later in the trial. Note, however, that it would not make sense to consider a case in which a unit undergoes scaling early in a trial and then exhibits absolute timing.

We next describe how to quantify these three schemes both at the level of the neural population and of individual neurons in RNNs trained on either 2-Context or 2-Stimulus tasks.

### Task structure differentially shapes the time encoding strategies at the population level

In order to visualize the internal dynamics of the RNNs we first plotted the normalized activity observed during the short and long intervals sorted according to the latency of peak activity for each unit during the short interval (Fig 4**A-B**, left panels), and sorted by the long intervals (Fig 4**A-B**, right panels). Interestingly, although the target output was a ramping pattern, relatively few RNN units appeared to be ramping. Rather, the global activity patterns in both tasks might be best conceptualized as neural sequences. Yet, while the self-sorted sequences appeared to be visually similar for both tasks, the cross-sorted sequences were dramatically different. Specifically, in the 2-Context task it appeared that neurons fired in the same order for both the short and long intervals—suggestive of a scaling encoding strategy. However, in the 2-Stimulus task the cross-sorted PSTHgrams revealed a more complex relationship between the spatio-temporal patterns of activity during the short and long intervals—suggestive of a more stimulus-specific encoding strategy.

**Fig. 4.**
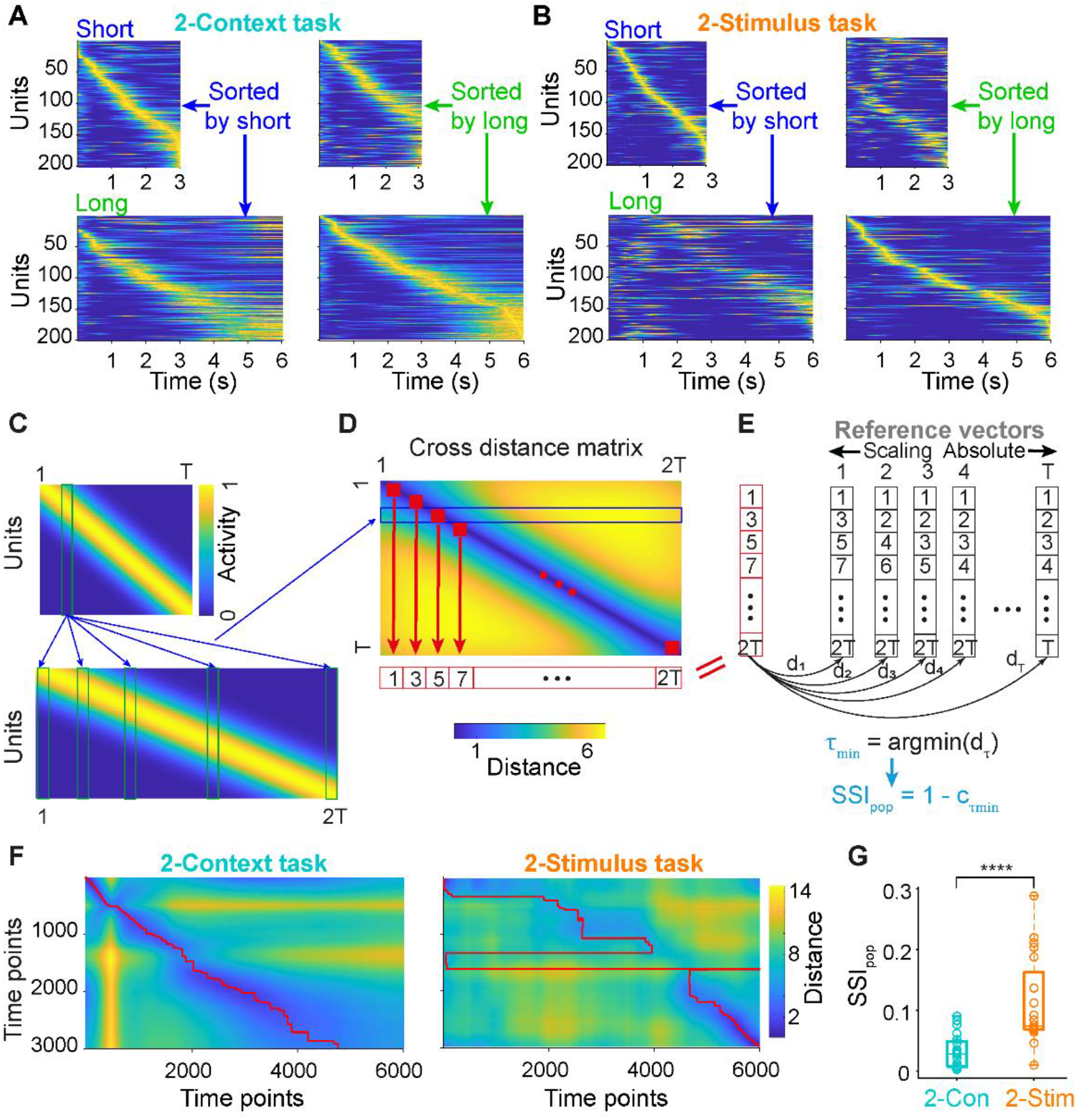
Distinct population dynamics in RNNs trained on the 2-Context and 2-Stimulus task. (**A**) Population activity for short (top) and long (bottom) intervals sorted according to the peak activity latency during short (left) and long (right) intervals for RNNs trained on the 2-Context task. (**B**) Same as **a** for the 2-Stimulus task. (**C**), (**D**), (**E**) Schematic of the calculation of the stimulus-specific index (SSI_pop_). A prototypical neural sequence that undergoes pure temporal scaling from the short (top) to long (bottom) intervals is used as an example (**C**). The vectors of the pairwise time points from the short and long dynamics are used to calculate all pairwise Euclidean distances, and these pairwise distances comprise the cross-distance matrix (**D**), in which a row (e.g., blue rectangle) represents the distances between one column vector of short dynamics and all column vectors during the long dynamics. The minimal index vector (red vectors in (**D**) and (**E**)) represents the indices along the x-axis that correspond to the minimal distances for each row of the cross-distance matrix (red squares). A series of reference vectors which vary from pure scaling to pure absolute timing (black vectors) are compared to the minimal index vector, and a value τ_min_ is defined as the τ at which the pairwise distance reaches the minimum. Finally, the correlation coefficient between the minimal index vector and the absolute-scaling reference vector at τ_min_ is used to calculate SSI_pop_. (**F**) Cross distance matrices for an example simulation of the 2-Context (left) and 2-Stimulus tasks (right). Red lines denote the indices of the minimum values for each row. (**G**) SSI_pop_ for RNNs trained on the 2-Stimulus task is significantly higher than that for 2-Context task (*n* = 20 simulations for each, two-sided Wilcoxon rank-sum test *P* < 0.0001). Boxplot: central lines, median; bottom and top edges, lower and upper quartiles; bottom and top whiskers: extremes.

To quantify if the neural dynamics observed in the 2-Context and 2-Stimulus tasks were more consistent with a scaling, absolute, or stimulus-specific code, we first developed as stimulus-specific index (SSI_pop_) based on previously described geometric approaches [12, 30, 31]. We started with the cross-Euclidean distance matrix between population dynamics for short and long intervals (see Methods), which compares the similarity of the activity across all time pairs during the short and long intervals (Fig 4**C-D**, example based on a case of perfect scaling of the entire population). We then extracted the index (time bin of the long interval) corresponding to the minimum value along each row of the cross-time distance matrix (red square in Fig 4**D**), which results in a vector of the time points that in the long-interval that are closest to each of the time points in the short-interval: the minimal index vector (red row vector in Fig 4**D** and column vector in Fig 4**E**). This minimal index vector was then matched to all possible reference vectors representing perfect scaling codes to a perfect absolute code (black column vectors in Fig 4**E**) by computing the distances d_τ_ between each pair (Fig 4**E**). The reference vector with the minimum distance (d_τmin_) to the minimal index vector denoted the best absolute-scaling vector. The correlation (c_τmin_) between the best absolute-scaling vector and the minimal index vector determines how good the match is: 1.0 reflecting perfect scaling, absolute timing, or a perfect mixture of absolute and scaling code. However, the correlation will be low or even negative in the case of a stimulus-specific code. Therefore, SSI_pop_ was defined by 1-c_τmin_ (Fig. 4**e**), meaning that both perfect scaling and absolute timing would result in an SSI_pop_=0, and the stimulus-specific code would be proportional to SSI_pop_.

We calculated SSI_pop_ for all 20 RNNs in both the 2-Context and 2-Stimulus tasks. SSI_pop_ was significantly higher during the neural dynamics of the 2-Stimulus task compared to the 2-Context task (Fig 4**G**), indicating that dynamics observed during the 2-Stimulus task reflected a stimulus-specific encoding strategy more so than the 2-Context task. However, consistent with the visual inspection of the dynamics and distance matrices (Fig 4**A, F**), it is clear that the 2-Stimulus task was not entirely accounted for by a stimulus-specific strategy, suggesting a mixed code. Thus we next examined the three encoding strategies from the perspective of the individual units in the network.

### Task structure shapes timing encoding strategy at the level of single units

To understand whether the encoding of the short and long intervals was most consistent with a scaling, absolute, or stimulus-specific code at the level of single units, we used a previously described measure of absolute-versus-scaling index (ASI) [23], and incorporated a novel stimulus-specific index (SSI_unit_) into the framework. Much as SSI_pop_ quantifies how different the dynamics of two neural populations are, SSI_unit_ quantifies how different the firing-rate profiles of a unit are during a short versus long trial (see Methods). More specifically, for a given unit, a high SSI_unit_ implies the temporal profiles during two trials are not related to each other through scaling or absolute timing. A low SSI_unit_ implies that the temporal profiles are related through scaling or absolute timing, thus justifying the use of the ASI to further quantify scaling versus absolute timing. To calculate the SSI_unit_ we first time-warped the temporal profile of a unit during the long interval into a series of reference absolute-scaling traces spanning from pure scaling to pure absolute timing with a mixture of both in between (Fig 5**A**). These reference traces were defined by a “breaking point” τ marking the transition from absolute timing to scaling (τ=0 reflects perfect scaling and τ =T_short_ reflect absolute timing). All reference traces were compared with the short dynamics by computing the Euclidean distance at each τ (d_τ_). The reference trace with the minimum distance (d_τmin_) denoted the best match with the actual temporal profile of the unit. Finally, as with SSI_pop_, the SSI_unit_ was defined as 1.0 minus the correlation between the temporal profile during the short intervals and the reference trace at τ_min_ (c_τmin_). For a given unit with a low SSI_unit_ (≤0.5), we went on to calculate its ASI which is also based on τ_min_ (see Methods). With the SSI_unit_ and ASI in hand, we classified a given unit as either a stimulus-specific unit (SSI_unit_>0.5), a scaling unit (SSI_unit_≤0.5, ASI≤0.5) or an absolute unit (SSI≤0.5, ASI>0.5) (Fig 5**B**).

**Fig. 5.**
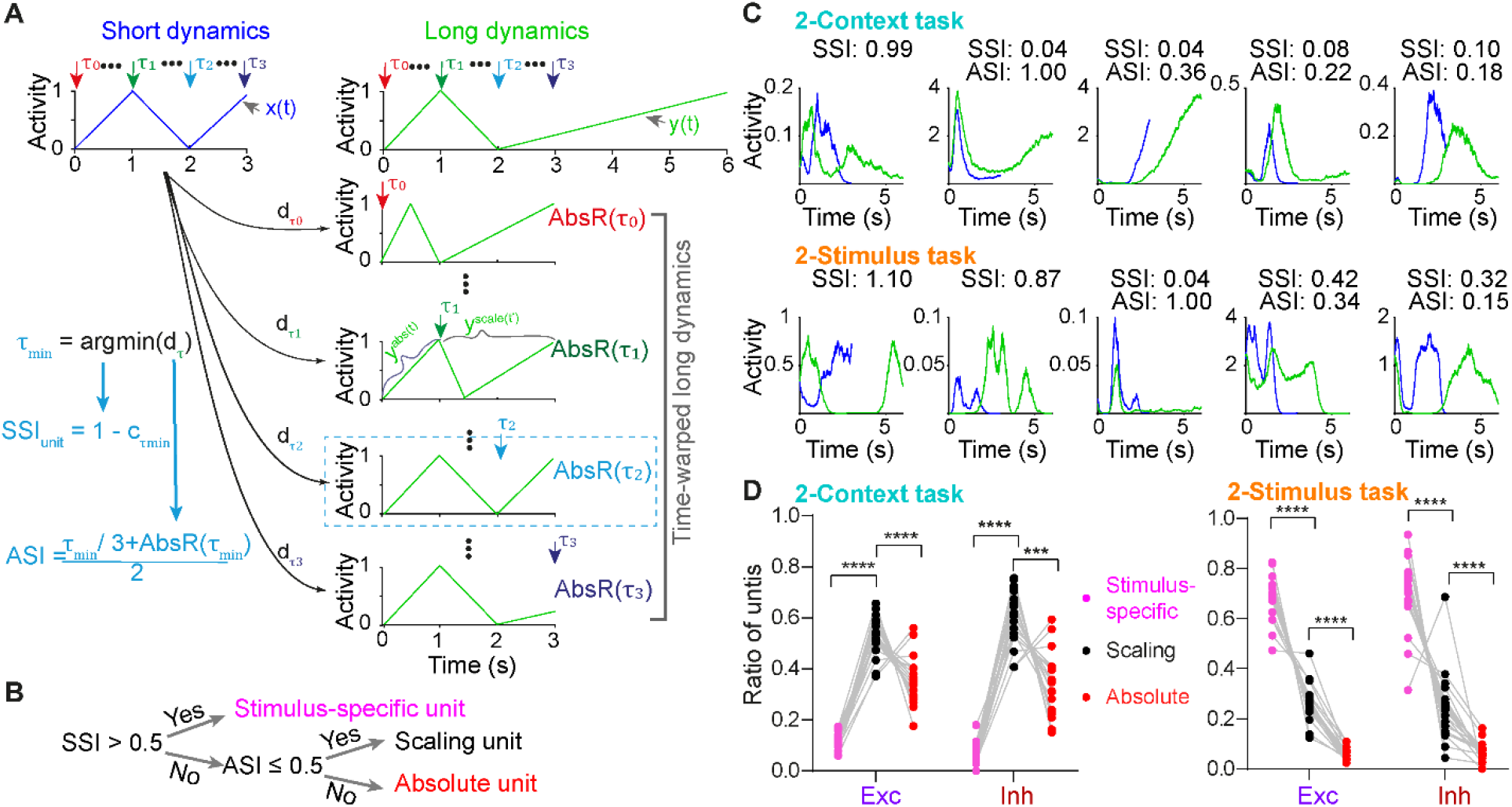
Different distribution of stimulus-specific, scaling, and absolute units between the 2-Context and 2-Stimulus tasks. **(A)** Schematic of the definitions of the stimulus-specific index (SSI_unit_) and absolute vs. scaling index (ASI) at the single unit level. Consider a hypothetical firing rate profile of a unit during a short (blue, x(t)) and long (green, y(t)) trial. As described in Methods, series of time-warped long dynamics are generated at breaking point τ_x_. before τ_x_ the dynamics are the same during both the short and long intervals (absolute timing, y^abs^(t)); after τ_x_ the dynamics is the scaled version of the corresponding original long dynamics (> τ_x_, scaling timing, y^scale^(t’)). Pairwise Euclidian distance between short dynamics and all time-warped long dynamics are computed at each τ_x_. The point at which the distance is minimal defines τ_min_ and is used to compute the SSI_unit_ as in SSI_pop_. To compute the ASI, a normalized measure of the distance before and after τ_min_ is calculated (AbsR) as in Methods. ASI is defined by τ_min_ and AbsR(τ_min_). (B) For a given unit, the SSI_unit_ is computed first, and if the SSI_unit_ is higher than 0.5, it is classified as stimulus-specific unit. If the SSI_unit_ is lower than 0.5, its ASI is computed, and it is classified as scaling unit if its ASI is lower than 0.5, otherwise as an absolute unit. (C) Dynamics of five example unit traces for short (blue) and long (green) intervals for the 2-Context (top) and 2-Stimulus (bottom) tasks, the corresponding SSI and ASI values are shown on top. Notice that for a given unit, ASI is only computed only when its SSI_unit_ is lower than 0.5. (D) For the 2-Context task (left), most units are classified as scaling units—for both excitatory and inhibitory units (n = 20 simulations, two-way ANOVA with repeated measures, for the unit classification factor : *F_(2,38)_* = 114.4 and *P* < 0.0001, posthoc Tukey tests *P* < 0.0001). For the 2-Stimulus task (right), stimulus-specific units are the most common (*n* = 20 simulations, two-way ANOVA with repeated measures, *F_(2,38)_* = 181.5 and *P* < 0.0001, posthoc Tukey tests *P* < 0.0001).

This approach allowed us to classify each unit of the network and contrast the distribution of temporal classifications between the 2-Context and 2-Stimulus tasks. These analyses revealed that RNNs exhibit a mixed encoding strategy, exhibiting a broad range of scaling, absolute, and stimulus-specific units (Fig 5**C**). However, there were highly significant differences in the distributions of temporal classes between the RNNs trained on the 2-Context and 2-Stimulus tasks (Fig 5**D**). The 2-Context RNNs were dominated by scaling units, while 2-Stimulus RNNs had more stimulus-specific units. The results partially explain why 2-Context RNNs were better at generalizing to novel intervals. Because our RNN structure obeyed Dale’s law it was possible to contrast the encoding strategies of excitatory and inhibitory neurons. Interestingly the distribution of scaling, absolute, and stimulus-specific cells appeared similar between excitatory and inhibitory neurons (Fig 5**D**).

To establish a causal relationship between the distribution of temporal classes to the functional properties of the RNNs we selectively deleted units of different classes from the RNNs trained on both tasks (S1 Fig 1**A**). We then investigated how the performance changed in response to these deletions. Performance and RMSEs across six deletion manipulations (stimulus-specific, scaling, and absolute temporal-classes for the excitatory and inhibitory populations) revealed inhibitory scaling units more severely impaired RNN function (S1 Fig 1**B, C**) for the 2-Context task. In contrast, no single manipulation condition more severely affected both performance and RMSE in the 2-Stimulus task (S1 Fig 1**D**, **E**). Somewhat surprisingly these results reveal that in the case of the 2-Context task a single subtype of inhibitory neurons—those that were classified as scaling units—are the most critical for network dynamics and encoding time. Whereas in the 2-stimulus task the coding strategy can be considered to be truly mixed, in the sense that all temporal classes and excitatory-inhibitory neurons seem to contribute more or less equally to the underlying dynamics and the encoding of time.

### Task structure differentially shapes the relationship between recurrent dynamics and input/output space

After quantifying how the different task structures shaped the encoding strategies, we sought to determine if the differences can be understood in terms of the relationship between RNN dynamics and the input/output subspaces. Generally, recurrent dynamics is driven by two sources: the interaction between the inputs and input weights, and between recurrent activity and recurrent weights. To start to understand how the inputs affected the recurrent dynamics and how the recurrent dynamics would lead to the output through the output weights, we first performed the principal component analysis on the concatenated dynamics of both intervals for each task (Fig 6**A, B**). We found that the first PCs explained more variance than that for the 2-Stimulus task (88.15±0.75% vs 69.72±0.73%, Fig 6**C**). We then projected the recurrent dynamics into the low dimensional space spanned by the first three PCs (Fig 6**A, B**). Visually in PC space, the dynamics of the two intervals for 2-Context task orbited close to each other, while that for the 2-Stimulus task formed two distinct trajectories—consistent with our findings that 2-Context task tended to use an absolute-scaling strategy while 2-Stimulus, a stimulus-specific strategy. These observations were further demonstrated by plotting the dynamics in response to generalization conditions (Fig 2). In the 2-Context task the dynamics across different inputs smoothly transitioned to nearby trajectories, while in the 2-Stimulus task the trajectories clustered around the two trained (short and long) trajectories (S1 Fig 2),.

**Fig. 6.**
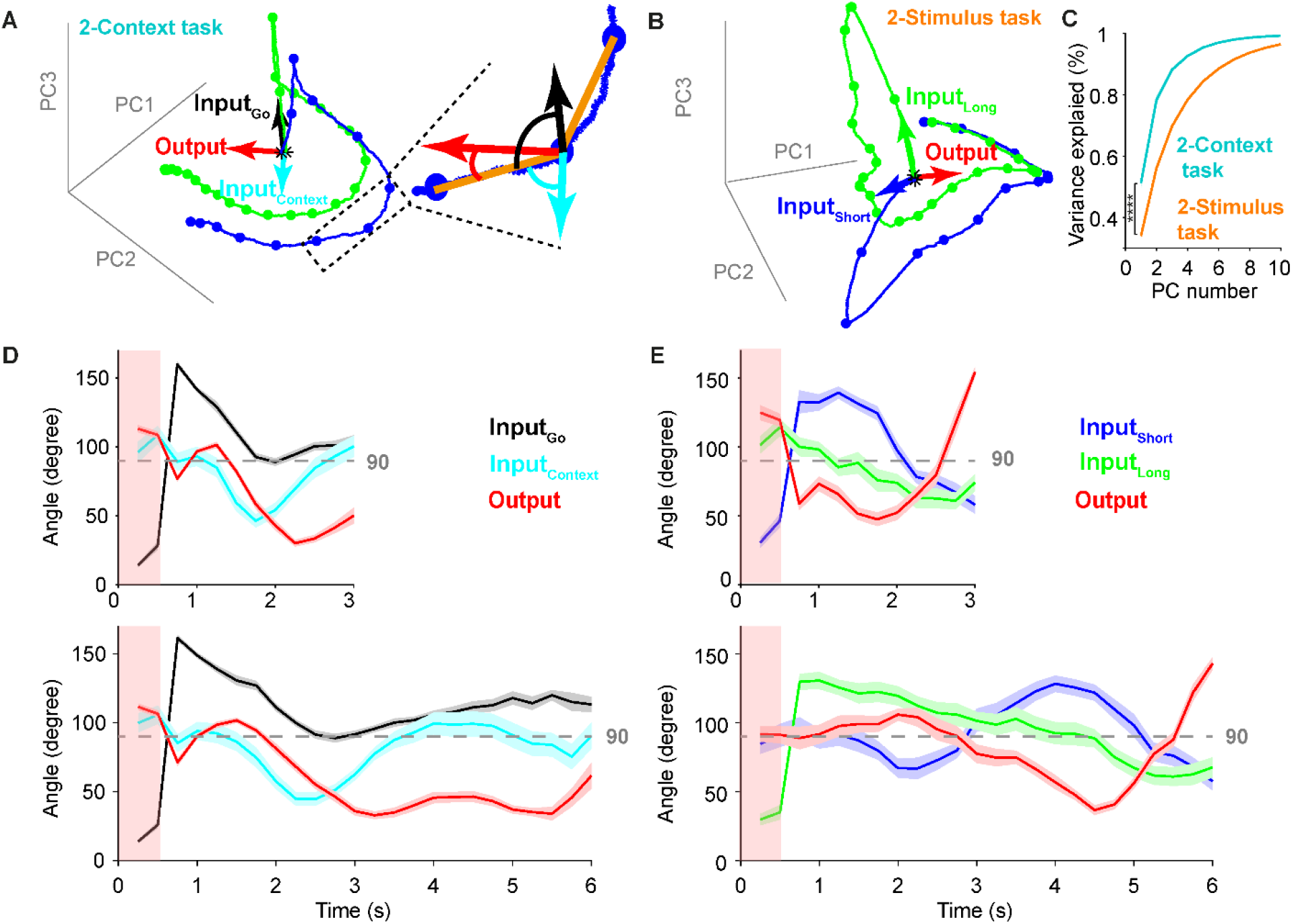
Differential subspace dynamics for RNNs trained on 2-Context and 2-Stimulus tasks. (**A**) For the 2-Context task, recurrent unit dynamics for the short (blue) and long (green) intervals were projected into the first three PC spaces. Astrisks denote the onset of inputs (t=0), arrows denote the corresponding weights vectors (Input_Go_, black; Input_Context_, cyan; and Output, red) projected onto the same PC space. Color dots denote the 250 ms intervals along each trajectory. Inset, schematic of angles between segments of the approximate RNN trajectory (orange) and the three weight vectors. These vectors were used to computed the pairwise angles to the Input_Go_, Input_Context_ and Output vectors. (**B**) Similar to **(A)** but for 2-Stimulus task, but here the two input vectors represented the Input_Short_ (blue) and Input_Long_ (green) weight vectors. (**C**) Same number of PCs explained more variance for 2-Context task than that for 2-Stimulus task (Two-way ANOVA, F_(1, 38)_ = 255.6 and *P* < 0.0001). (**D**) Average pairwise angles between segments of short (top)/long (bottom) dynamics and inputs/output vectors as in **(A)** for 2-Context task (20 simulations, data presents as Mean ± SEM). Shaded area donted the duration of the transient Input_Go_ **(E)** Same as in **(D)** but for 2-Stimulus task. Shaded area donted the duration of the transient Input_Short_ and Input_Long_.

To directly compare the relationship between the recurrent dynamics across time and the input/output weights, we projected the input weights—Input_Go_ and Input_Context_ for the 2-Context task, Input_Short_ and Input_Long_ for the 2-Stimulus task—and the output weights into the same PC space. We then computed the pairwise angles between the projected input/output vectors and each segment vector of recurrent dynamics across time (see Methods) (Fig 6**A**) for both tasks. Interestingly, for the 2-Context task the dynamics of both intervals first evolved in the Input_Go_ input direction as revealed by the small angle for the first 2 segments. After that, both trajectories stayed in a plane almost orthogonal to the Go input till the end of the trial. The dynamics were almost orthogonal to the Input_Context_ at the beginning (with angles close to 90 degrees) and then the angle decreased in the middle period and increased again to about 90 degrees at the later period. Finally, for output weights, the angle stayed close to 90 degrees at the beginning then it decreased to a low level till the end of the trial indicating that the dynamics followed the output weights directions in the later period of the trials to better generate the target ramp staring at the middle point of each trial.

For the 2-Stimulus task, the dynamics of short and long intervals started to follow their corresponding input directions and then went to the opposite directions after input offset and stayed almost orthogonal thereafter. While for the output weights, the angle started at around 90 degrees and then decreased around the start point of the target ramp then it increased at the end of the trials to the opposite direction.

### Task structure differentially shapes the learned recurrent synaptic connectivity

Ultimately the task-specific differences in RNN dynamics must be attributed to differences in input structure and the recurrent weight matrix. Thus we next characterized the relationship between the recurrent weight matrices and performance. Since our RNNs respected Dale’s law, we grouped weights into the four standard subtypes: all excitatory to excitatory unit connections (E→E), all excitatory to inhibitory unit connections (E→I), all inhibitory to excitatory unit connection (I→E), and all inhibitory to inhibitory unit connections (I→I). We then completely deleted each group of synapses and quantified the change in output performance (Fig 7**A**). Interestingly, deleting all E→E connections only slightly affected the performance and RMSE for both tasks, while deleting all other three groups decreased the performance or increased the RMSE. Deleting the I→E connections produced the largest change in RMSE (Fig 7**B**). We next quantified the connection probability and mean weights of each group (Fig 7**C**). Consistent with the performance and RMSE results, I→E connections exhibited the highest connection probability and mean weights for both tasks. Interestingly, to achieve similar output performance, the two tasks seemed to rely on different strategies in the structural level: 2-Context task favored higher connection probability, while 2-Stimulus task preferred higher mean weights (Fig 7**C**).

**Fig. 7.**
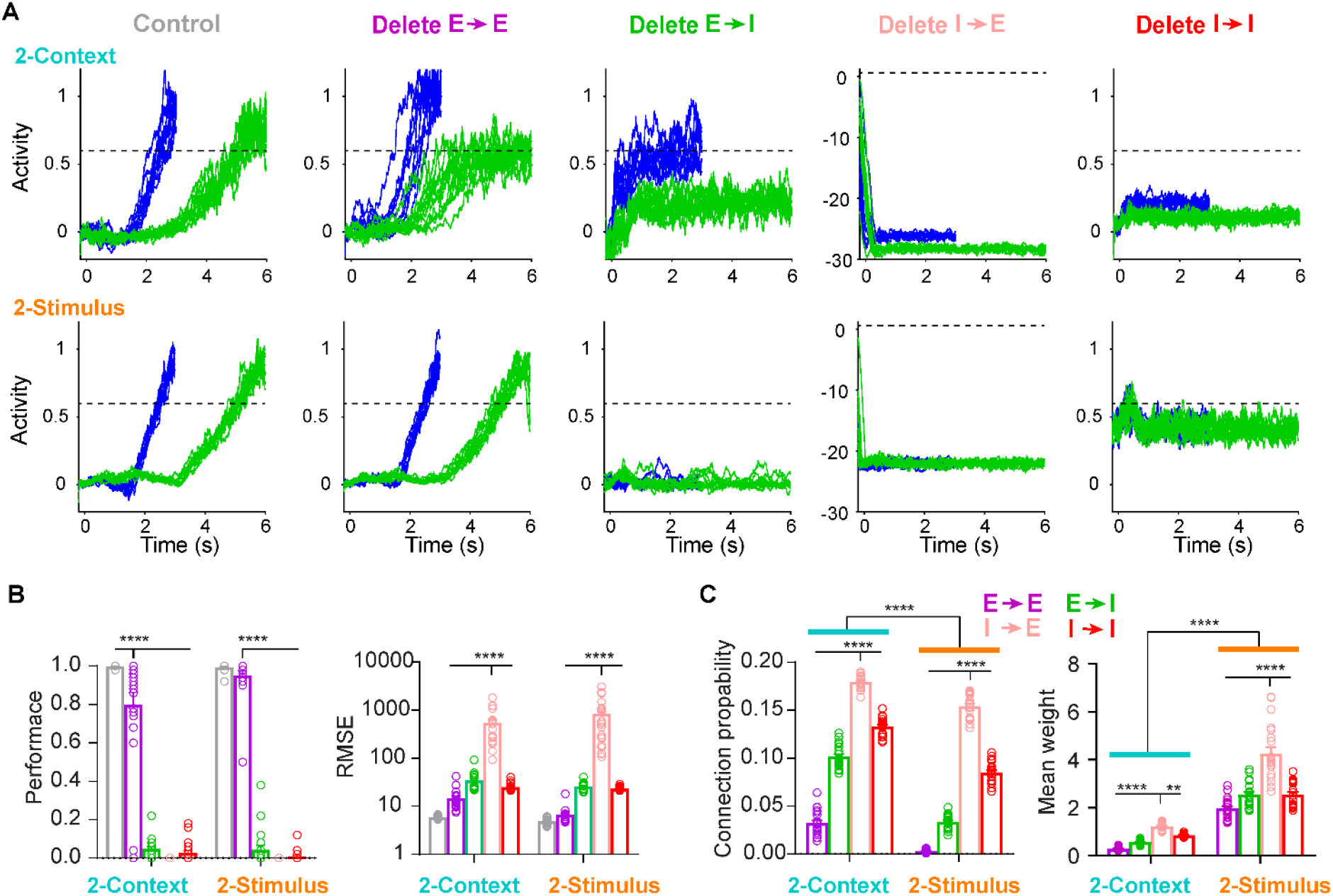
Differential connectivity patterns in RNNs trained on the 2-Context and 2-Stimulus tasks. (**A**) Example of the effects of deleting entire subgroups of synpases on performance in the 2-Context (top) and 2-Stimulus (bottom) tasks. From left to right, example output traces of the short (blue) and long (green) intervals for control condition, and after deleting all excitatory unit to excitroy unit connections (Delete E→E), all excitatory unit to inhibitory unit connections (Delete E→I), all inhibitory to excitatory unit connections (Delete I→E), and all inhibitory unit to inhibitory unit connections (Delete I→I). **(B)** Mean performance (left) and RMSE (right) of the outputs corresponding to the conditions in panel **A**. The performance for Delete E→E condition is significantly lower than the control but signigicanly higher than the other conditions in 2-Context task. For the 2-Stimulus task performance for Delete E→E was not significantly worse than the control, but significantly higher than the other conditions (two-way ANOVA with mixed-effect design, *F_4,152_* = 823.9, *P* < 0.0001, posthoc Tukey tests *P* < 0.0001). The RMSE for Delete I→E condition is significantly higher than the other conditions in both 2-Context and 2-Stimulus task (two-way ANOVA with mixed-effect design, *F_4,152_* = 39.8, *P* < 0.0001, posthoc Tukey tests *P* < 0.0001). **(C)** Left, connection propability in the 2-Context task was significantly higher than in the 2-Stimulus task(two-way ANOVA with mixed-effect design, *F_1,38_* = 338.3, *P* < 0.0001 for the task factor). Propability for the I→E connctions is significantly higher than that for the other three conditions: E→E, E→I, I→I in both 2-Context and 2-Stimulus task (*F_3,114_* = 2884, *P* < 0.0001 for the connection factor, posthoc Tukey tests *P* < 0.0001). Right, mean weight in the 2-Context task is significantly lower than that in the 2-Stimulus task (two-way ANOVA with mixed-effect design, *F_1,38_* = 219.1, *P* < 0.0001 for the task factor). Propability for the I→E connctions is significangly higher than that for the other three conditions: E→E, E→I, I→I in both 2-Context and 2-Stimulus task (*F_3,114_* = 183.7, *P* < 0.0001 for the connection factor, posthoc Tukey tests). ****=P < 0.0001, and **=P = 0.002.

### RNNs trained for the 2-Stimulus task are more robust to noise

We have seen that RNNs trained for the 2-Context task are better suited for generalization to novel intervals and this feature is related to the underlying dynamics being governed by a scaling encoding scheme. A question that emerges from these results is whether there is a computational tradeoff between the distinct dynamic regimes observed in both tasks? For example, does the pronounced stimulus-specific encoding in the 2-Stimulus task offer any potential advantages other than the categorical timing. As a first step to address this question we analyzed the robustness of both tasks in response to noise. In the brain, of course, neural networks are continuously subject to extraneous noise, and thus robustness to noise imposes an important constraint on biologically functional dynamic regimes [32]

As above we first trained RNNs on either the 2-Context and 2-Stimulus tasks with the standard settings, namely noise level of 0.45 (σ in Eq. 1), then we tested the networks by applying different values of σ. Example output traces for the 2-Stimulus task under all noise levels tested were less scattered than that for the 2-Context (Fig 8**A**). This was supported by the fact that the mean RMSE for the 2-Stimulus task was lower than that for the 2-Context (Fig. 8**B**). For both tasks, at high noise levels, there were some incorrect trials (< 10% and no significant difference between the two tasks) in which either the output never crossed the threshold during the trial or crossed the threshold outside of the acceptance windows We then directly contrasted the temporal precision of the correct trials and found that the standard deviations for the 2-Stimulus task were lower than that for the 2-Context task (Fig. 8**C**). Taken together, we conclude that the dynamic regimes underlying timing in the predominately stimulus-specific dynamics that emerged in the 2-Stimulus task provided a computational benefit in terms of robustness to noise. suggesting computational tradeoffs between different dynamic regimes for the encoding of time.

**Fig. 8.**
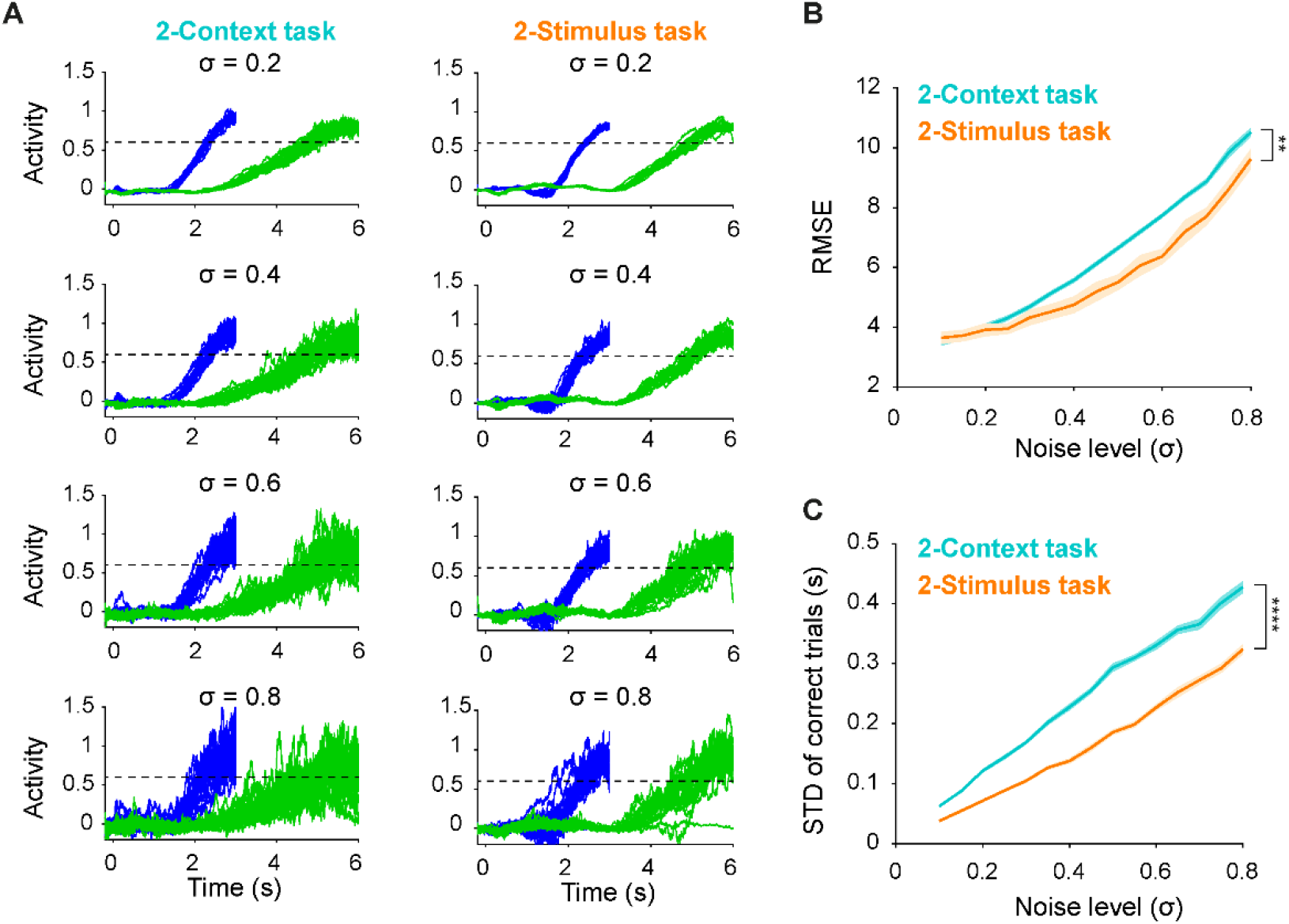
RNNs trained on the 2-Stimulus task were less sensitive to noise perturbations. (**A**) Output traces for short (blue) and long (green) intervals from an example RNN trained on the 2-Context (left) and 2-Stimulus (right) across different levels of noise (σ) during testing. (**B**) Mean RMSE (across 50 trials) for 2-Context task (cyan) is higher than that for 2-Stimulus task (orange) (n = 20 simulations, two-way ANOVA with mixed-effect design, *F_1,38_* = 9.35, *P* = 0.004). (**C**) Mean standard deviation of the time of threshold-crossing across all correct trials for 2-Context task (cyan) is higher than that for 2-Stimulus task (orange) (*F_1,38_* = 341, *P* < 0.0001). Data are presented as mean ± SEM.

## Discussion

Here we trained supervised RNNs on two simple temporal tasks that required the production of identical temporal output patterns based on previous behavioral results [23]: a ramping increase in output firing rate that peaked after either a short (3 s) or long (6 s) interval. The tasks differed only in how the short and long intervals were cued: either by a continuously presented context input (2-Context task) or by two distinct brief inputs (2-Stimulus task). In principle the same dynamic regimes could have emerged and solved both tasks, yet, significantly different dynamic regimes emerged in the different tasks. Thus depending on the task RNNs encoded time in different ways, and exhibited fundamentally different computational properties, particularly regarding how the networks generalized to novel stimuli.

### Neural dynamic regimes of population clocks

A converging body of experimental and computational evidence suggests that neural circuits encode time in spatiotemporal patterns of neural activity. Two experimentally relevant neural dynamics regimes by which neurons can encode time include ramping activity and population clocks. Ramping codes generally refer to monotonically increasing (or decreasing) firing rates throughout an interval [24, 33–39]—in ramping codes firing rate often peaks at the time of the target interval, and in principle, a single neuron can encode time throughout the entire duration. Population clocks refer to time-varying patterns of activity in which time is encoded in the population activity of neurons, which generally exhibit nonmonotonic changes in firing rate, and importantly these dynamics are generated by the recurrent connectivity within a neural circuit [1, 11, 40, 41]. Population clocks can include either simple sparse neural sequences as well as complex spatiotemporal patterns in which a given neuron can exhibit multiple timing fields [28, 42–50].

In the current simulations, the target output patterns were a simple ramping pattern, yet most of the units in the RNNs were not well described as ramping units—even though it seems that this would be the simplest and most direct solution to solve the tasks. Rather, the neural dynamics observed in the RNNs studied here, are most consistent with the notion of population clocks in general and neural sequences in particular (Fig 4). These results are in line with other computational models in which neural sequences encode time [51–54]. The reason RNNs trained with supervised learning rules seem to converge to neural sequences rather than ramping activity are not well understood, but it has been recently proposed that neural sequences represent a fairly optimal encoding scheme for downstream neuron (the output unit in our case) to read out time [23].

### Absolute, scaling, and stimulus-specific codes

We outlined three general temporal encoding strategies by which a population of neurons could solve temporal tasks that require producing multiple intervals (Fig 3)—such as the two tasks examined here. The scaling strategy is perhaps the most intuitive because it essentially exploits the same neural dynamics to produce both a short or long interval by altering the speed at which the dynamics unfold. Indeed, such scaling has been observed experimentally [14, 16, 23, 26, 38]. Neurons that exhibit absolute timing have also been experimentally observed [14, 23, 26, 27, 55–57], as have stimulus-specific codes in which the same or different intervals can be encoded in different neural trajectories [17, 46, 55, 58–60]. To date, however, these different encoding strategies have not been carefully analyzed or quantified. To this end, we described two general purpose quantitative measures—the ASI and SSI_unit_—that can be applied across a wide range of single-unit data and used to classify neural responses.

These measures revealed a different distribution of unit types across the RNNs trained on the 2-Context and 2-Stimulus tasks (Fig 5). Specifically, over 50% of the units in the 2-Context RNNs were classified as scaling units, whereas in the 2-Stimulus RNNs over 50% were classified as stimulus-specific units—that is, their temporal profiles between the short and long interval were not consistent with either absolute or scaling coding strategies. This differential distribution is consistent with the intuition that because in the 2-Context task the context input is active during both the short and long intervals, and a stimulus-specific encoding strategy is more difficult to implement compared to the 2-Stimulus task—i.e., the input space of the 2-Context task is smaller. Put another way, in the 2-Stimulus task RNNs are likely to begin their trajectories at the beginning of each trial (t=0) in more distant regions of neural state space than in the 2-Stimulus task.

The differential distribution of scaling, absolute, and stimulus-specific neurons accounts in part for the distinct computational features of both types of networks. Specifically, the classification of units into different temporal coding strategies allowed us to demonstrate that selectively deleting some classes impaired RNN performance more than others. Deleting a few scaling units impaired RNN performance in the 2-Context task significantly more than deleting absolute or stimulus-selective units. In contrast in the 2-Stimulus task, all classes contributed to performance with an approximately equal weighting—reflecting a much more mixed encoding strategy [61, 62].

### Computational trade-offs between time-encoding dynamic regimes

The 2-Context and 2-Stimulus tasks required producing the same temporal output patterns but generated dramatically different behaviors when challenged with novel inputs. Of particular relevance was that in response to novel levels of activation of the inputs, the 2-Context RNN exhibited a smooth scaling of the temporal profile of the output. In this task, in response to the *go* stimulus, RNN’s generated a neural trajectory that resembled a neural sequence. Depending on the analog value of the context input this trajectory unfolded at either a slow or fast speed to produce the short or long interval, respectively. Critically, in response to novel levels of activation of the tonic context input the velocity of the neural trajectory varied smoothly—thus generating smooth temporal scaling of the output pattern. This same property has been observed in numerous other models of timing [12, 30, 34, 63–65]. Specifically, a single input or variable is able to modulate the velocity of the RNN dynamics in an approximately linear fashion.

In contrast to the temporal scaling behavior observed in the RNN trained on the 2-Context task, when the 2-Stimulus RNNs were tested with inputs they were not trained on (e.g., 50% Input 1 + 50% Input 2) they did not exhibit smooth generalization. Importantly, they also did not exhibit catastrophic degradation—i.e., the internal dynamics was robust to very different initial states. Rather they exhibited categorical timing—essentially a winner-take-all competition between two distinct trajectories. This result is consistent with the notion that RNNs can encode multiple neural trajectories in regimes that have been referred to as dynamic attractors [31, 66], locally stable transient trajectories [67], or stable heteroclinic channels [68, 69]. Here, two trajectories serve as basins of attraction (or more accurately “rivers-of-attraction”) which lead the activity of the network into one or the other of the two dynamic attractors.

Both temporal scaling and categorical timing are behaviorally relevant examples of timing. Specifically, many tasks require smoothly scaling the temporal output patterns or categorically discriminating or producing one of distinct two intervals [12–15, 21, 22]. Thus, we have shown that the population clocks that emerge in RNNs can account for both temporal scaling and categorical timing and that it is possible to distinguish between both regimes based on the percentage of units that undergo scaling or stimulus-specific timing.

It is also relevant to note that RNNs learned to solve the 2-Context task in fewer training trials than the 2-Stimulus task. This may be because it is easier to adjust weights to generate a single trajectory at two different speeds than to generate largely distinct trajectories. Furthermore, during training the 2-Context task the RNN is always subject to tonic external input which in effect might facilitate learning by suppressing the potential emergence of chaotic regimes [70, 71].

### Experimental predictions

As is evident from the behavioral data, a wide range of distinct neural regimes, from ramping activity to a diverse range of neural population clocks, have been observed experimentally across different brain areas and behavioral tasks [for reviews see: 1, 6, 7, 10]. Here we show that the same is true even in RNNs trained on two tasks that require the production of the same temporal output patterns. Our results thus suggest that much of the experimentally observed variability might be accounted for by relatively subtle differences in task structure. Furthermore, because most timing tasks used in laboratories tap into ecologically relevant behaviors, different tasks may encourage generalization patterns that best approximate their ecological relevance. These distinct generalization patterns will, in turn, result in time being encoded in different dynamic regimes—e.g., regimes that are well-suited for temporal scaling or categorical timing.

A number of strong experimental predictions emerge from our results. First, at the behavioral level, we predict that whether rodents are trained on the 2-Context or 2-Stimulus will lead to different generalization patterns to novel stimuli. For example, a single odor along with a tone context stimulus could be used for the 2-Context task, and two brief odors as the stimuli in the 2-Stimulus task. We predict that changing the loudness of the tone in the 2-Context task will scale the output pattern, but mixing the odors will result in categorical timing rather than the production of an intermediary interval. Second, we predict that neural recordings from animals trained on these tasks will exhibit specific neural dynamic signatures, i.e., in the 2-Context task more neurons will be categorized as scaling units compared to the 2-Stimulus task. Of course, one must take into account that results may be dependent on the brain areas being recorded. However, based on the current literature we expect this prediction to hold in those areas that have been implicated in timing across many tasks, including the striatum, supplementary/secondary motor areas, and prefrontal cortical areas.

## Materials and Methods

### Firing-rate RNN model

RNNs were based on firing-rate units that obeyed Dale’s law (N = 200, 80/20% excitatory/inhibitory). RNN dynamics was described by the following equations:

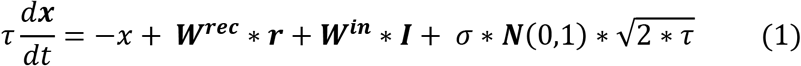

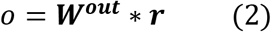

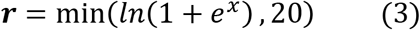

where 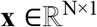 represents the input currents of RNN units, and firing rate vector **r** is obtained by applying a Softplus function constrained by an upper bound of 20. The time constant τ was equal to 100 ms for all units. 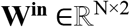 and **I** are the input weights and external inputs, which are task-specific as described below. Each unit received independent Gaussian noise **N**(0,1) with the standard deviation of 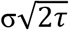. Unless otherwise specified, σ = 0.45. 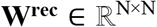 is the recurrent weight matrix. Self-connections were absent in the network. The output (*o*) of the network is computed linearly from the output weights **W^out^** and **r**. RNNs were implemented and trained in Tensorflow starting from the code of Kim et al [72].

#### Training

Networks were trained using adaptive moment estimation stochastic gradient descent algorithm (Adam) to minimize the root mean square error loss (RMSE) between network output o and target z:

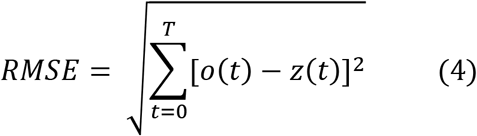

where T is the total length of a given trial. The target and mask are task-dependent as described below. The learning rate was 0.01, and other TensorFlow default values were used.

Only recurrent weights **W^rec^** and output weights **W^out^** were trained. **W^rec^** was initialized as a random sparse matrix with a connection probability of 0.2. **W^rec^** was initialized from a normal distribution and transformed to absolute values. To begin in an approximately balanced regime the inhibitory weights were multiplied by 4. To respect Dale’s law during training a rectified linear operation on **W^rec^**, and excitation and inhibition were implemented by multiplication with a diagonal matrix of 1 and −1 representing excitatory and inhibitory units, respectively [73]. **W^out^** was initially drawn from a standard normal distribution. **W^in^** was also drawn from a standard normal distribution and was fixed during training.

During training, a discretization step of 20 ms was used. After training, RNNs were ported to Matlab using the trained parameters and a discretization step of 1ms.

Parameters were updated every trial. After Every 100 trials of training, the network was tested for 100 trials to compute the task performance (see below) and mean RMSE. When task performance was higher than 97% and the mean RMSE is lower than 2, the training was considered a success and stopped

### Interval tasks

#### 2-Context task

Inspired by the timing task used by Wang et al [14] in which context cues indicated the lengths of intervals, we designed a 2-Context two-interval task. In this task, the output of the RNN needs to generate either a short (3 s) or long (6 s) interval in each trial. For a given training trial with length *T*, two external inputs 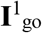 and 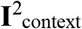 were applied at stim_onset_ after a baseline with random durations between 0.2 s ms and 0.6 s. Specifically,

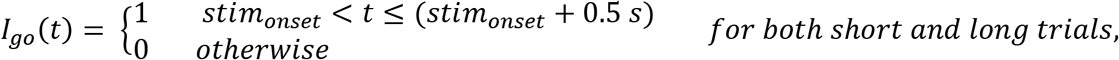

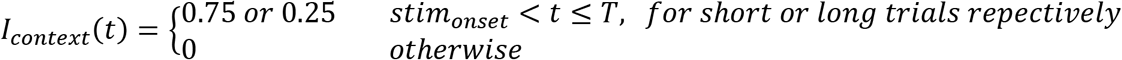

The output targets were defined as:

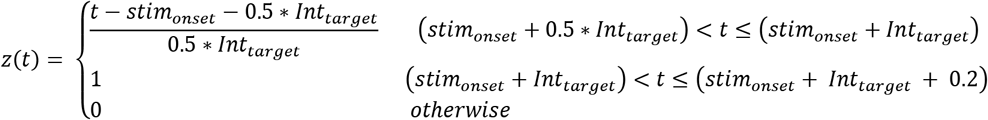

where the target intervals (Int_target_) were 3 and 6 seconds for the short and long trials, respectively.

#### 2-Stimulus task

The 2-Stimulus task was based on a two-interval odor discrimination task [23], which required the production of the identical output patterns as the 2-Context task. However, the short and long intervals were cued by two different inputs **I**_short_ and **I**_long_ which like the **I**_go_ in the 2-Context task stepped up from 0 to 1 for a brief 0.5 s period.

#### Task performance

Response time for a given trial was defined as the time when the output crosses a threshold of 0.6. The correct trials were defined as those in which the output crossed the threshold within an acceptance window between stim_onset_ + 0.5 Int_target_ and stim_onset_ + Int_target_. Task performance was defined as the ratio of correct trials among all testing trials.

Unless otherwise specified, the “delay” epoch (stim_onset_ to stim_onset_ + Int_target_) was used for analysis.

### Generalization to novel inputs

To test how the RNN trained on the 2-Context task would generalize to novel intervals, we first trained the RNN using normal setting up for the 2-Context task, namely **I**_context_ of 0.75 and 0.25 for the short and long trials, respectively. Then we tested the trained RNNs by gradually varying the context level from 0.75 to 0.25 with steps of 0.05. Fifty trials of each level were obtained for analyses.

After training in the 2-Stimulus task generalization to novel inputs was tested by gradually varying the ratio of **I**_short_ and **I**_lon*g*_ with steps of 0.1 so that the sum of both inputs was always 1.

#### Correlation measure

To quantify changes in the temporal profile of the output units across different inputs during generalization tests we first computed the correlation coefficient between the mean response times (when the output crosses the threshold) and the generalization conditions for both the 2-Context and 2-Stimulus tasks (the absolute values of the correlations were used due to the negative correlation for the 2-Context task).

#### Sigmoid slope measure

To further quantify generalization to novel inputs in both tasks we also fitted the mean response times to the input conditions with a sigmoid function as follow:

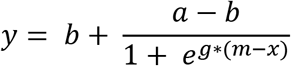

Standard nonlinear least square methods implemented in Matlab were used to optimize the fits. We then compared the slope *g* for both tasks. Higher *g* values reflect more categorical generalization.

### Prototypical dynamical regimes for timing two intervals

To illustrate the possible neural dynamical strategies used for timing two intervals—scaling, absolute, and stimulus-specific, we generated three pairs of prototypical dynamics for the short (3 s) and long (6 s) intervals composed of 100 units with the time step of 0.001 s. In such settings, the dynamics for the short and long interval were represented as 100×3000 and 100×6000 matrices respectively, with the row being units and column being time points.

The dynamics for long interval were the same for all three strategies, which was described as:

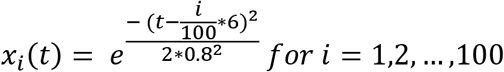

where dynamics of all units were Gaussian functions with the same variance but different means uniformly spanned the whole 6 s. The dynamics for the short interval were different for the three strategies and were defined as follows:

#### Scaling

The dynamics for the short interval in the scaling strategy was simply a matrix of uniform subsampling of the time dimension of the long dynamics.

#### Absolute

For the absolute strategy, the dynamics of the first 50 units for the short interval were the same as that for the long interval.

#### Stimulus-specific

For stimulus-specific example, we first uniformly subsampled the time dimension of the long dynamics matrix to 3 s. Then we randomized the order of the unit indices.

### The stimulus-specific index at the population-level (SSI_pop_)

To quantify how well the short and long neural trajectories can be explained by the stimulus-specific strategy at the population level, we developed a novel stimulus-specific index in population-level (SSI_pop_), which is largely based on establishing that the trajectories are not consistent with temporal scaling or absolute timing. We first obtained the mean population dynamics (Δt=1 ms) for two intervals by averaging across 25 trials, which led to two matrices, **X_short_** (200×3000) and **X_long_** (200×6000). We computed the pairwise Euclidean distance between **X_short_** and **X_long_**, which led to the distance matrix **D** (3000×6000). We then obtained the index of the minimum values across each row of **D**, which led to the minimal distance vector **I_min_** (3000× 1), which partially captures the relationship between the population dynamics for the short and long intervals.

Next **I_min_** was contrasted with a reference matrix **R** (3000×3000) with τ indexing the column:

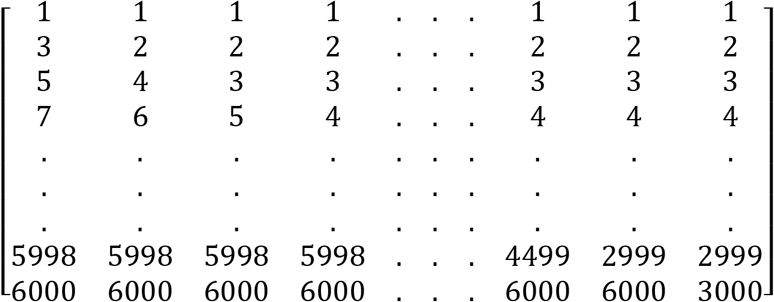

Specifically, a given column vector corresponding to τ in **R** is defined as:

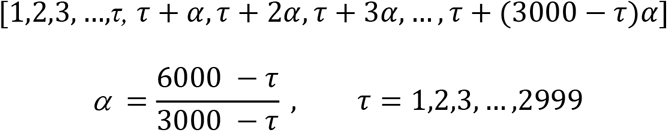

Each column vector (3000× 1) in **R** represents one absolute-scaling reference profile spanning from pure scaling (τ = 1) to pure absolute (τ = 3000), with mixed profiles in between in which absolute timing transitions to scaling at τ with the scaling factor α varied to keep the length of each vector the same. We then computed the Euclidian distances between **I_min_** and all the column vectors of **R** and extracted the vector with the minimum distance at τ_min_, which indicates the best reference vector that can be used to explain the **I_min_**. Note that the construction of the R matrix accounts for units that fire throughout the entire trial—thus capturing the properties of a neuron that always fired at the end of the trial (e.g., a potential motor neuron). It is also possible to build R by fixing the scaling factor at 2 after each point τ, in which case the last element of each column in R would progressively change from 6000 to 3000. We have run analyses with this partial scaling approach as well with qualitatively similar results.

Finally, the SSI_pop_ was defined as:

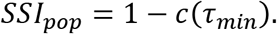

where the c(τ_min_) is the correlation between the **I_min_** and reference vector at τ_min_. For pure scaling dynamics for the two intervals as an example, **I_min_** should be the main diagonal of distance matrix **D**, [1, 3, 5, 7, …, 6000], which makes τ_min_= 1, corresponding to the pure scaling reference vector. Consequently, the c(τ_min_) is 1 and SSI_pop_ is 0. That indicates that the pure scaling dynamics can not be explained by stimulus-specific strategy but by absolute-scaling strategies, more specifically, the scaling strategy.

### Stimulus-specific index and absolute-scaling index (ASI) for single units

We extended a previous description of an absolute vs. scaling index (ASI) for single units [23], by including a novel measure of the stimulus-specific profile: the stimulus-specific index at the single-cell level (SSI_unit_ as in Fig 5**A**). As described previously we searched for the best transformation of dynamics for the long interval (y(t)) to that for the short interval (x(t)), by concatenating an absolute portion of the long response (y^abs^(t)) and a temporally scaled portion of the long response scaled by a factor α (y^scale^(t’)). More specifically, we searched for a breakpoint τ to divide y(t) into an absolute and scaled segment, that best matches x(t), as measured by the Euclidean distance (Dist(τ)). Specifically,

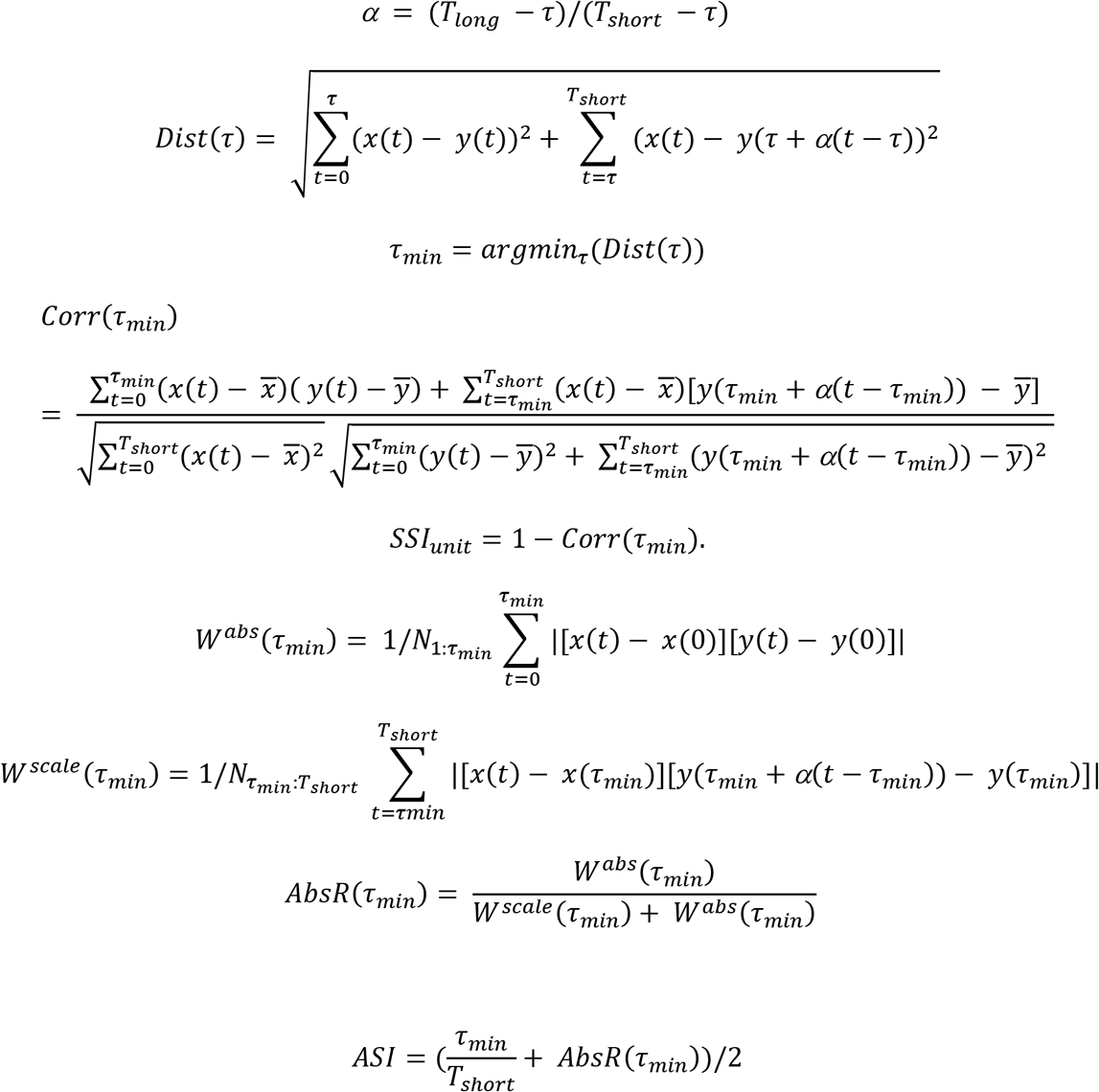

τ spans all possible breakpoints from 0 to T_short_ (for the short interval and T_long_ for the long interval). The segment before τ denotes the absolute period and the period after τ denotes the segment scaled by α for the long response. τ_min_ corresponds to the breakpoint with the minimal Euclidian distance Dist(τ_min_). Different from previous work, we also computed the correlation coefficient between x(t) and transformed y(t), Corr (τ_min_). Then the SSI_unit_ is defined as that 1 minus Corr(τ_min_). In the following steps, the absolute and scaling weights are calculated between dynamics for the short interval and the time-warped dynamics for the long interval at τ_min_ with *N_a:b_* being the number of time points between a and b, and absolute ratio AbsR(τ_min_) was also calculated. The absolute temporal factor corresponds to τ_min_ /T_short_, and ASI was defined as the average of the absolute temporal factor and the AbsR (τ_min_).

To classify each unit according as a stimulus-specific, scaling, or absolute unit we first calculated SSI_unit_ for each unit. We then classified a unit as stimulus-specific if SSI_unit_ was > 0.5; if the SSI_unit_ was ≤ 0.5 looked at its ASI and classified it as is an absolute unit if ASI > 0.5, or as a scaling unit if ASI ≤ 0.5.

### Unit-deletion and weight deletions experiments

Based on the classification of units being stimulus-specific, scaling, or absolute, we ran deletion experiments to start to understand the causal role of each type of unit. For a given unit to be deleted, we removed all the connections attached to that unit in connection matrix **W^rec^** as in Eq. 1 and then ran the RNN with the rest parameters fixed. We tested various numbers of deleted units in each type. For a given condition, we randomly selected the deleted cells from the pool 10 times and repeated each deletion experiment for 20 trials for each interval. Then performance and RMSE were averaged across all selections and trials.

To quantify how much each class of connection types—E→E, E→I, I→E and I→I connections—contributed to the recurrent dynamics and output performance, we performed synapse deletion experiments. Similar to the unit deletions, for a specific class of connections, we set all the weights of that group to be zeros while leaving the other weights unchanged. Performance and RMSE were then computed for each condition.

### Pairwise angle analysis

To understand the relationships between the RNNs trained on 2-Context and 2-Stimulus task and the input/output subspace (Fig. 6) defined by the inputs weights and output weights, we first performed principal component analysis (PCA) on the concatenated mean dynamics for the short and long intervals. We then projected the original dynamics into the first three PCs. We then binned the projected dynamics into segments of 250 ms. For a given segment, a vector was obtained by subtracting its start point from its end point. Finally, we computed the pairwise angles between all such segment vectors across time and input/output weight vectors (in the same PC space of the original input/output weight vectors).

### Noise perturbation experiments

To test the robustness of the outputs of the RNN trained on the 2-Context and 2-Stimulus tasks, we first trained the two tasks with noise level σ = 0.45 as in equation (1). We then tested the trained RNNs with various levels σ from 0.1 to 0.8 for 50 trials for each interval. We then compared the RMSE between the outputs and targets for all trials and the standard deviation of the crossing times for the correct trials. Note that for all conditions tested, the incorrect trials were less than 10% for both tasks, and there was no significant difference for that between the two tasks.

### Statistical analyses

Statistical analyses were carried out with standard functions in MATLAB (MathWorks) and Prism (GraphPad Software). The sample size, type of test, P values, and the F values for ANOVA are indicated in the figure legends. All data and error bars represent the mean and SEM except for the boxplot in Fig. 4, where median and quartiles were presented. In all figures, the convention is *: P < 0.05, **: P < 0.01, ***: P < 0.001, ****: P < 0.0001.

## Author contributions

Conceptualization: SZ, DVB

Methodology: SZ, DVB

Simulations: SZ

Analysis: SZ, DVB

Supervision: SCM, DVB

Writing—original draft: SZ, DVB

Writing—review & editing: SZ, SCM, DVB

## Data and codes availability

All data are available in the main text or supplementary materials. Codes used for all simulations in this paper are available at (https://github.com/ShanglinZhou/RNN_2Intervals). Codes used for producing the figures in this work will be also available at the same place.

## Supporting information

**S1 Fig.**
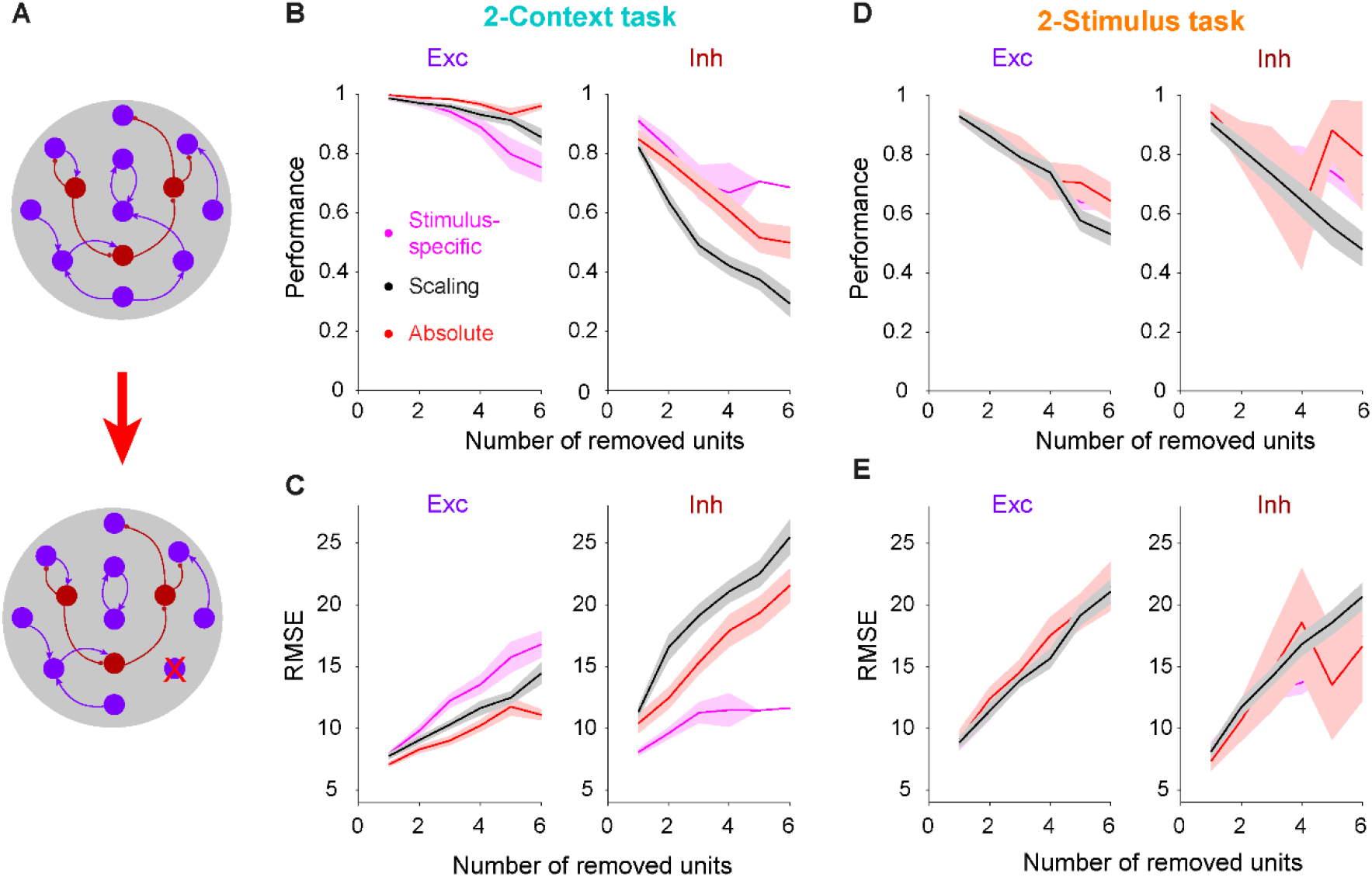
Differential functional effects of deleting specific classes of units. (**A**) Schematic of the deletion experiments. To delete a given unit denoted by the red arrow (bottom), all in and out weights of the recurrent weight matrix of that units were set to zero. (**B**) Performance of RNNs trained on the 2-Context task after progressively deleting units from specific temporal-classes: stimulus-specific, scaling, and absolute temporal-classes for both excitatory (left) and inhibitory (right) units. For each data point, units were randomly selected 10 times, and 10 test trials obtained. A three-way ANOVA revealed highly significant effects of main temporal-class (*F_2,619_* = 31, *P* < 10^−12^) and Ex-Inh (*F_2,619_* = 390, *P* < 10^−66^) factors. Additionally, there was highly significant interaction between temporal-class and Ex-Inh class (*F_2,619_* = 27, *P* < 10^−10^) and multi-comparison analyses showed that performance for inhibitory scaling cells was significantly lower than all other 5 deletion manipulations (P < 0.0001 for all comparisons). (**C**) Similar to (**B**) but for RMSE. As in (**B**), there were highly significant main effects (*F_2,619_* = 34, *P* < 10^−14^, and *F_2,619_* = 118, *P* < 10^−24^, for temporal-class and Ex-Inh, respectively), as well as a significant interaction between temporal-class and Ex-Inh (*F_2,619_* = 46, *P* < 10^−18^). And again the inhibitory scaling cells increased the RMSE more than all other deletion manipulations (*P* < 0.0001 for all comparisons). (**D-E**) There were no main effects of temporal-class or Ex-Inh that were consistently significant for both the performance and RMSE measure. The interaction between temporal-class and Ex-Inh was either trending (*F_2,619_* = 2.5, *P* =0.08) or mildly signficant (*F_2,619_* = 3.6, *P* = 0.027) for the performance and RMSE analyses, respectively. Data are presented as performance mean ± SEM across 20 RNNs.

**S2 Fig.**
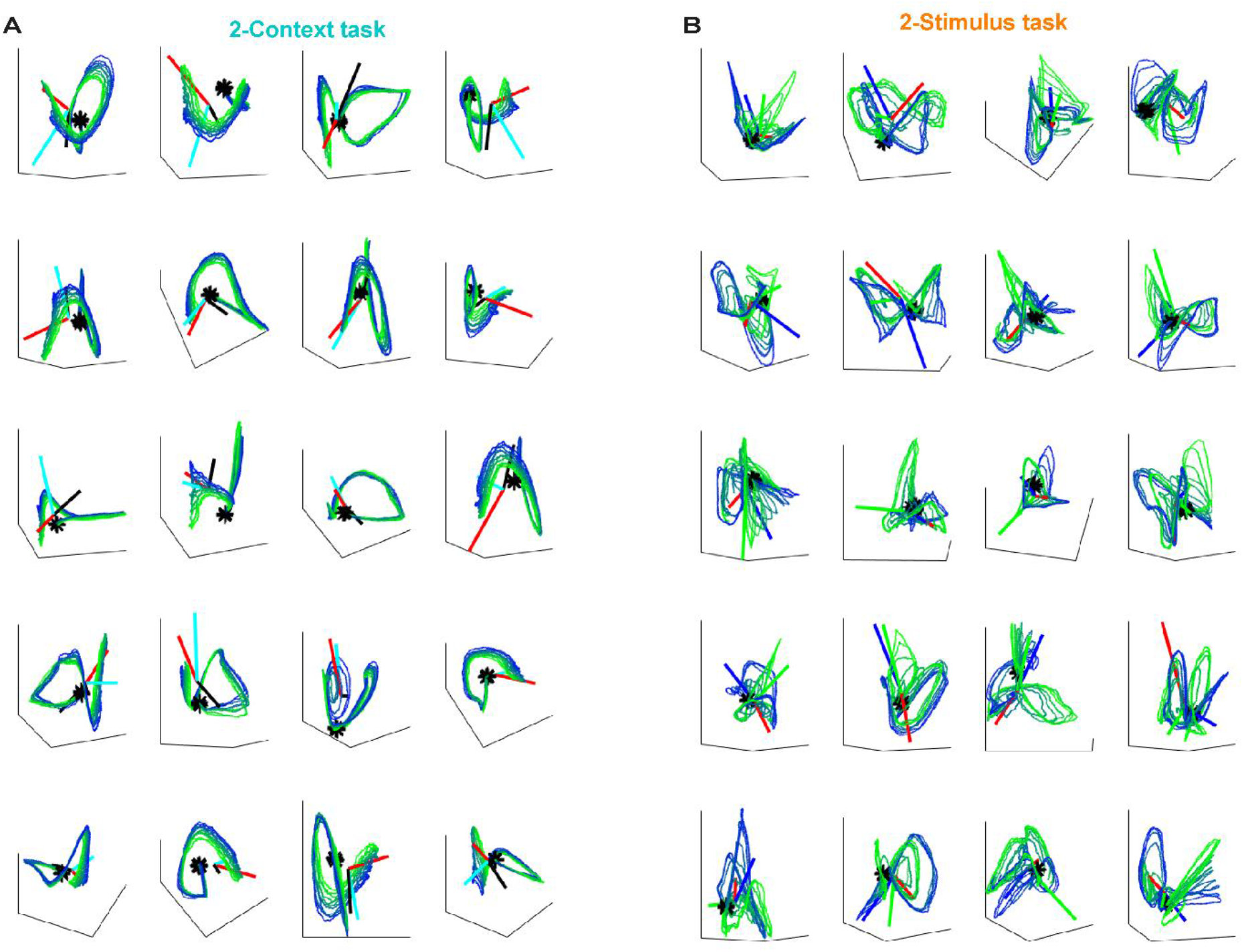
PCA plots of the recurrent dynamics for generalization to novel intervals. (**A**) Recurrent dynamics corresponding to different context level (denoted by the color) as in Fig. 2 were projected into the first three PCs in 20 RNNS trained on 2-Context task. The arrows denoted the directions of Input_Go_ (black), Input_Context_ (cyan) and Output (red) weights projected into the same PC space. (**B**) similar as in **(A)** but for the 2-Stimulus task.

## Notes

### Competing Interest Statement

The authors have declared no competing interest.

